# A statistical approach for identifying primary substrates of ZSWIM8-mediated microRNA degradation in small-RNA sequencing data

**DOI:** 10.1101/2022.02.17.480958

**Authors:** Peter Y. Wang, David P. Bartel

## Abstract

**Background:** One strategy for identifying targets of a regulatory factor is to perturb the factor and use high-throughput RNA sequencing to examine the consequences. However, distinguishing direct targets from secondary effects and experimental noise can be challenging when confounding signal is present in the background at varying levels.

**Results:** Here, we present a statistical modeling strategy to identify microRNAs (miRNAs) that are primary substrates of target-directed miRNA degradation (TDMD) mediated by ZSWIM8. This method uses a bi-beta-uniform mixture (BBUM) model to separate primary from background signal components, leveraging the expectation that primary signal is restricted to upregulation and not downregulation upon loss of ZSWIM8. The BBUM model strategy retained the apparent sensitivity and specificity of the previous ad hoc approach but was more robust against outliers, achieved a more consistent stringency, and could be performed using a single cutoff of false discovery rate (FDR).

**Conclusions:** We developed the BBUM model, a robust statistical modeling strategy to account for background secondary signal in differential expression data. It performed well for identifying primary substrates of TDMD and should be useful for other applications in which the primary regulatory targets are only upregulated or only downregulated. The BBUM model, FDR-correction algorithm, and significance-testing methods are available as an R package at https://github.com/wyppeter/bbum.

## INTRODUCTION

Differential expression (DE) analyses seek to identify gene products that change in abundance after either altering a condition or perturbing a regulatory factor. Aiding in these analyses are statistical pipelines, such as DESeq2 [1] and edgeR [2], which compare RNA-seq or microarray datasets to identify RNAs with significantly changed levels, after correcting for false discovery rate (FDR) due to multiple testing using the Benjamini-Hochberg procedure [3]. These pipelines have been invaluable for DE analyses of mRNAs as well as noncoding RNAs.

Noncoding RNAs often subject to DE analyses include the microRNAs (miRNAs), which are small RNAs that direct widespread post-transcriptional repression of metazoan mRNAs [4]. To perform this function, miRNAs associate with the effector protein Argonaute (AGO) to form a complex in which the miRNA specifies which targets are repressed, primarily through base pairing between the seed of the miRNA (miRNA nucleotides 2–7) and a site in the target mRNA [5]. Meanwhile, AGO causes repression, typically by recruiting the cytoplasmic mRNA deadenylation machinery [6].

Most miRNAs are quite stable, with half-lives much greater than those of typical mRNAs, presumably a consequence of their association with AGO, which protects miRNAs from cellular nucleases [7, 8]. However, some miRNAs are more rapidly degraded, and in cells of both mammals and flies, this more rapid degradation appears to be the result of target-directed miRNA degradation (TDMD) [9]. TDMD is a phenomenon in which targets with unusual complementarity cause a conformational change recognized by the ZSWIM8 E3 ubiquitin ligase, which polyubiquitinates the AGO protein, leading to its degradation by the proteasome, thereby exposing the miRNA to degradation by cellular nucleases [9, 10]. Through DE analysis of small-RNA sequencing data acquired after reducing ZSWIM8 in different cell types, more than 40 candidate miRNA substrates of ZSWIM8 have been identified [9].

When identifying candidate substrates of ZSWIM8, standard DE analysis is not sufficient to distinguish between miRNAs that are significantly upregulated due to the primary effect of losing ZSWIM8-mediated TDMD, and those with significantly perturbed expression due to secondary effects, such as transcriptome changes caused by the dysregulation of miRNAs or other changes that might be caused by the loss of ZSWIM8. To exclude miRNAs changing due to secondary effects, Shi et al. [9] use the knowledge that ZSWIM8 mediates degradation of miRNA substrates, which implies that these substrates should undergo only upregulation upon the loss of ZSWIM8. Accordingly, the significance cutoffs (*α* values) of FDR-adjusted *p* values from DESeq2 (*p*_*adj*_) are each adjusted down to the most permissive level that excludes all downregulated miRNAs.

As a result, these ad hoc adjustments of *α* values vary widely, ranging from 0.05 to 10^−7^ for different datasets analyzed (Fig. 1A) [9]. Although this approach seems better than classifying any significantly upregulated miRNA as a ZSWIM8 substrate, it has several shortcomings: 1) it is unduly sensitive to outliers among downregulated miRNAs, which can reduce sensitivity; 2) FDRs are inconsistent among experiments and cannot be predetermined; and 3) the FDR and specificity of each analysis depend on the sample size.

**Figure 1.**
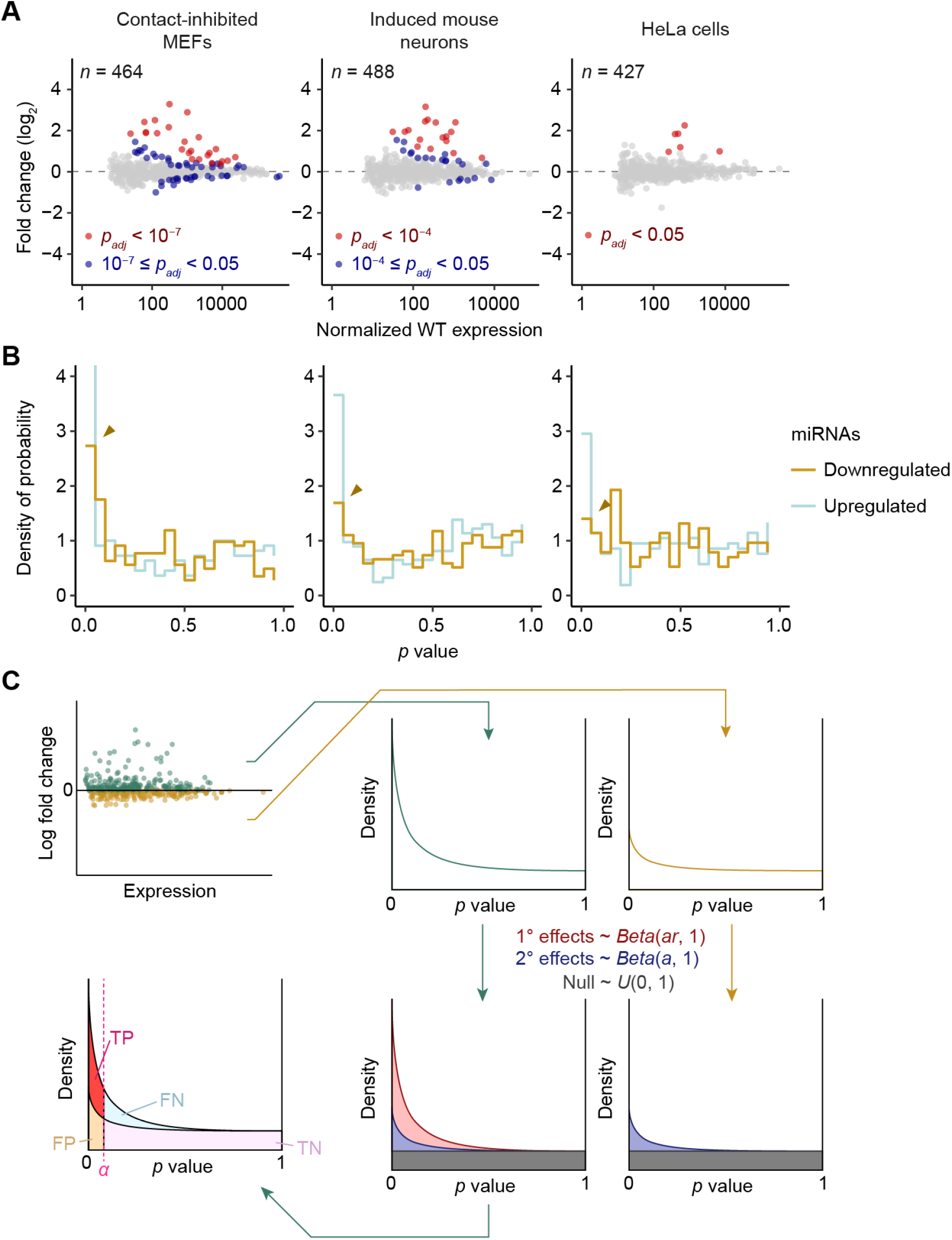
Statistical modeling of secondary effects in DE datasets. **(A)** Representative plots of fold changes in miRNA levels observed upon *ZSWIM8* knockout, as measured by small-RNA sequencing. Points for miRNAs with significant upregulation based on the cutoffs defined by Shi et al. [9] are colored in red. Points for miRNAs that did not meet the adjusted cutoffs but would be significant based on a common cutoff of 0.05 are colored in blue. The number of miRNAs and passenger strands quantified in each dataset is indicated. **(B)** Histograms of raw *p* values for upregulated and downregulated miRNAs and passenger strands analyzed in datasets of **A**. The peak near *p* = 0, which indicates true signal above null, is indicated by an orange caret for downregulated miRNAs in each dataset. **(C)** BBUM modeling of *p* values from DE datasets. *p* values corresponding to upregulated and downregulated miRNAs are fit in parallel to the BBUM model in which downregulated points are fit to a distribution missing the beta component for primary effects. Based on the fitted model, expected FDR and other statistics can be calculated for any cutoff *α* (TP, true positives; FN, false negatives; FP, false positives; TN, true negatives).

Here, we developed a statistical modeling strategy that accounts for varying secondary effects in RNA-seq datasets, thereby enabling primary substrates to be identified at a consistent predetermined statistical stringency. Compared to the previous approach, this strategy achieved more robust results with fewer shortcomings. This strategy should also provide an improved strategy for identifying direct targets of other types of regulatory pathways.

## RESULTS

### Significant signal among downregulated miRNAs implies secondary effects

In published datasets, a standard *p*_*adj*_ cutoff at *α* = 0.05 was suitable for identifying primary substrates in some contexts, such as HeLa cells [9]. However, in other contexts, such as contact-inhibited mouse embryonic fibroblasts (MEFs) or induced mouse neurons (iMNs), the same cutoff would have classified many miRNAs with changes that seemed no different than background as primary substrates (Fig. 1A). These differences observed between datasets, which motivated the use of varying stringency of *α* values, were presumably due to varying levels of secondary effects. To evaluate this idea, we examined the downregulated miRNAs, reasoning that because loss of ZSWIM8 should only cause upregulation of primary miRNA substrates, any signal among downregulated miRNAs in excess of that expected by chance implied the existence of true secondary effects. To search for this signal, we examined *p*-value distributions. A set of data points drawn from a null distribution is expected to have a uniform distribution of *p* values, ranging from 0 to 1, as the cumulative fraction of points called as false positives should equal to *α* for all values of *α*. Thus, a significant signal should manifest as a peak of enriched density near *p* = 0 [11]. For each of the three datasets of Figure 1A, the distribution of raw *p* values from DESeq2 was examined, looking separately at the results for upregulated and downregulated miRNAs. As expected for datasets that included ZSWIM8 substrates, distributions for upregulated miRNAs peaked near *p* = 0 (Fig. 1B). In addition, for the two datasets that required a more stringent *α* value, the distributions for downregulated miRNAs also peaked near *p* = 0, albeit at a level lower than that observed for upregulated miRNAs (Fig. 1A, B). These results supported the idea that some miRNAs were truly downregulated upon the loss of ZSWIM8, likely as a result of secondary effects, and contexts with stronger secondary effects required stronger adjustments of stringency.

### Statistical modeling of *p* values enables the separation of primary and secondary effects

If secondary effects acted symmetrically, causing miRNAs to increase as well as decrease (at similar frequency and similar magnitudes), then the excess in the peak near *p* = 0 observed for upregulated miRNAs compared to that observed for downregulated miRNAs should correspond to the density of true primary substrates of ZSWIM8 (Fig. 1C). Accordingly, we developed a statistical strategy to separate the components of the *p*-value distribution to better classify the primary substrates and the secondary effects. Our strategy made three assumptions: 1) primary effects were stronger than secondary effects, 2) secondary effects were approximately symmetrical between upregulated and downregulated data points, and 3) primary effects caused upregulation and never downregulation. Previous studies have described the successful use of a beta-uniform mixture (BUM) distribution model and its variants to model *p*-value distributions between 0 and 1 [12–14]. In these studies, the uniform component represents the distribution of null data points, while the beta component represents the characteristic peak near *p* = 0. Building upon these concepts, we implemented a modified mixture distribution model, which we call the bi-beta-uniform mixture (BBUM) model, to describe our *p* values. The BBUM distribution contains two beta components [*Beta*(*ar*, 1) and *Beta*(*a*, 1)], instead of one, which respectively correspond to the primary and secondary effects, followed by a similar uniform component [*U*(0, 1)] for the null distribution (Fig. 1C). The *p* values were fit to this mixture model, with the downregulated data points fit to a model that lacked the primary-effect component (Fig. 1C).

The model faithfully captured the distributions of *p* values from both halves of each dataset, especially for the *p*-value density near 0 (Fig. 2). As expected, datasets that required more stringent *α* values for *p*_*adj*_ (Fig. 1A) were modeled with more pronounced beta components for secondary effects, as indicated by the greater deviation between the null distribution and the model fit for downregulated points (Fig. 2, orange shading). These results indicated that for these datasets the model was able to separate primary and secondary effects.

**Figure 2.**
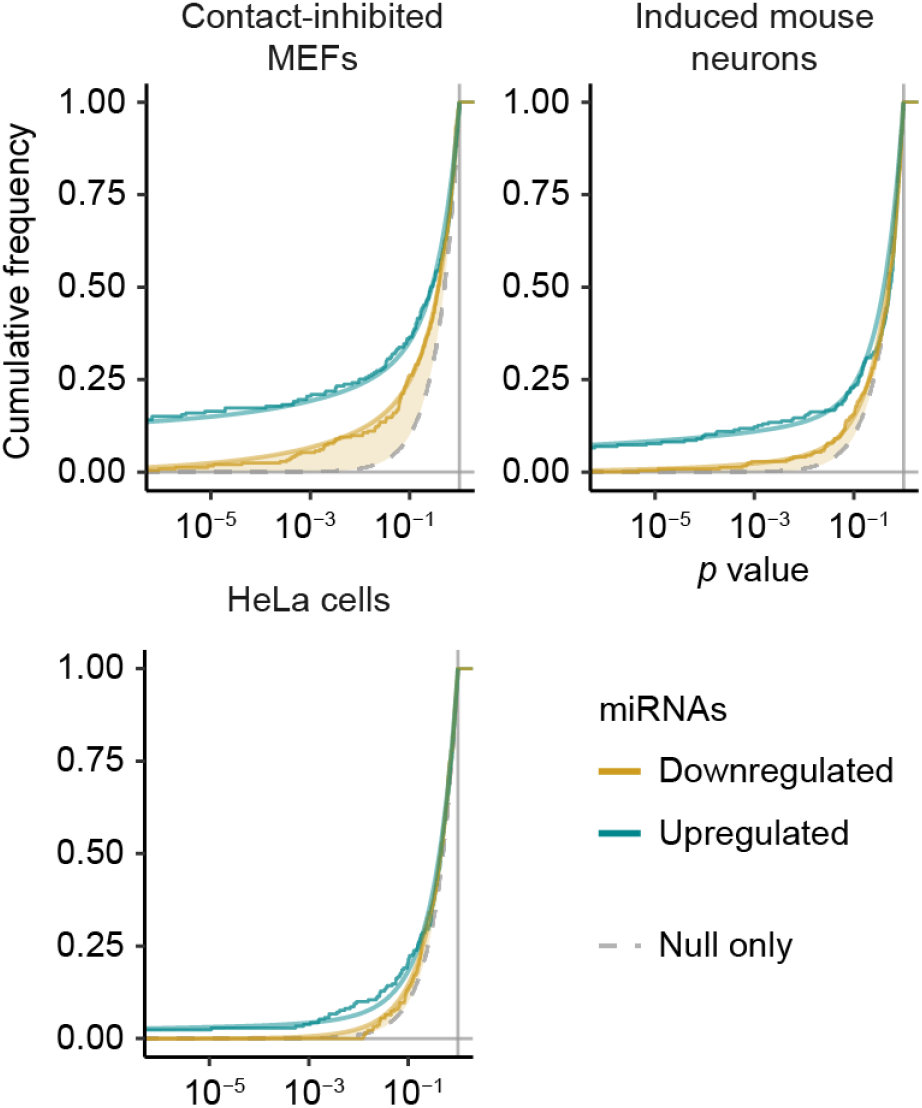
Fit of empirical *p* values to the BBUM model. Plotted for each dataset are empirical cumulative distributions of *p* values for upregulated miRNAs (turquoise) and downregulated miRNAs (orange), with the respective cumulative distributions of the fitted BBUM model overlaid as smooth lines. The uniform null distribution is shown as a gray dashed line. Deviation between the distribution for downregulated miRNAs and the null distribution is shaded in light orange; a greater deviation at the lower tail indicates a greater excess in density near *p* = 0, which corresponds to a more substantial contribution of the beta component for secondary effects in the BBUM model.

### Significance testing after BBUM adjustment predicts direct substrates of TDMD

Because the model was able to represent the distribution density that corresponded specifically to the primary effects, the expected FDR could be calculated at any desired cutoff among the *p* values considered, which we defined as the BBUM-FDR-adjusted *p* value (*p*_*BBUM*_). Choosing a *p*_*BBUM*_ cutoff of 0.05, we reanalyzed the datasets from mammalian and fly cell lines from Shi et al. [9] to identify ZSWIM8 primary substrates. Across all nine datasets examined, the proposed primary substrates identified using the BBUM strategy largely matched those identified previously, while imposing a consistent, predetermined FDR cutoff (Fig. 3A). Out of the 75 instances classified as significant at this cutoff, four were newly classified as significant. Three of these four involved miRNAs that were either also proposed to be ZSWIM8 substrates in other contexts or related to another proposed ZSWIM8 substrate, which supported the idea that these four miRNAs included true ZSWIM8 substrates (Fig. 3A, Table S1). This idea was further reinforced by analysis of the passenger strands of these candidate miRNAs. During miRNA biogenesis, the pre-miRNA hairpin is processed into a miRNA duplex containing the miRNA paired to its passenger strand. When this duplex associates with AGO, the miRNA strand is retained, and the passenger strand is ejected and rapidly degraded. Because TDMD acts upon mature AGO−miRNA complexes, the miRNA strand and not the passenger strand should be affected by the loss of ZSWIM8 [9]. Indeed, each of the four newly significant miRNAs were upregulated upon the loss of ZSWIM8 without significant change in their passenger strands, as observed for other miRNAs predicted to be ZSWIM8 substrates (Fig. 3B).

**Figure 3.**
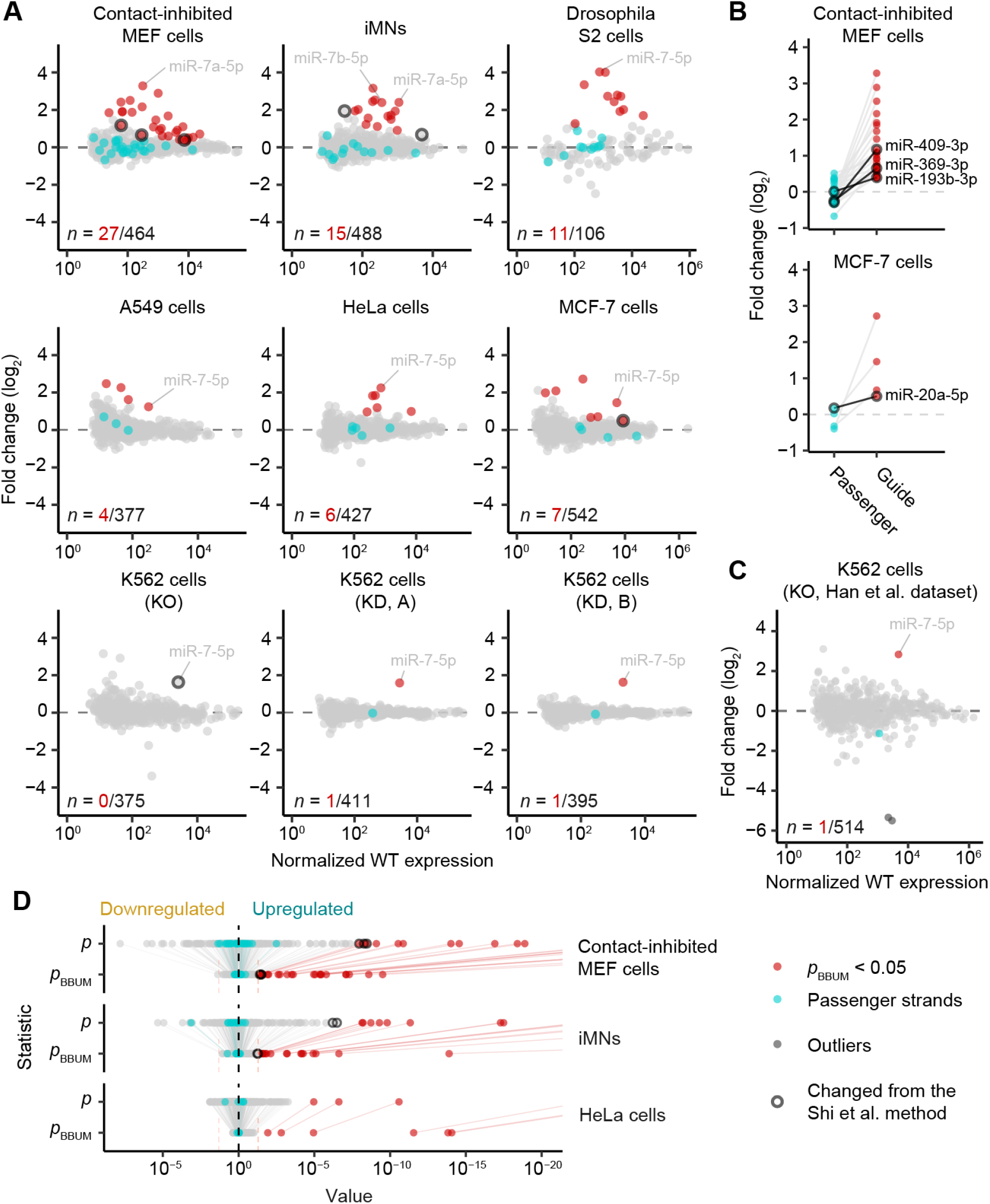
Identification of candidate ZSWIM8-sensitive miRNAs using the BBUM method. **(A)** Plots of fold changes in miRNA levels observed upon loss of ZSWIM8 in all mammalian and fly cell lines examined in Shi et al. [9]. Two datasets were derived from CRISPRi knockdown (KD) of *ZSWIM8* in K562 cells, using one of two different guide RNAs (A or B). All other datasets were derived from knockout (KO) of either *ZSWIM8* or *Dora*, the Drosophila *ZSWIM8* ortholog. Points for miRNAs that met the common *p*_*BBUM*_ significance cutoff of 0.05 are in red, with *n* indicating the number passing this cutoff as fraction of the total number of miRNAs and passenger strands quantified. Points for passenger strands of these miRNAs are in cyan, if the passenger strands were quantified. Points for miRNAs with classifications differing from that of previous work [9] are outlined in black (*n* = 7). Points for miR-7-5p, a known TDMD substrate [15], are labeled. **(B)** Different effects of the loss of ZSWIM8 on miRNAs and their passenger strands. Fold changes in miRNA levels are shown for two datasets with newly significant miRNAs. Only miRNAs found significant by the BBUM method and whose passenger strands were quantified in the dataset are shown, with each miRNA paired with its passenger strand(s). Points for newly significant miRNAs are outlined and labeled. **(C)** Plot of fold changes in miRNA levels observed upon knockout of *ZSWIM8* in K562 cells by Han et al. [10]. Points for two downregulated outliers (hsa-miR-221-3p, hsa-miR-222-3p) are shown in darker gray. Otherwise this panel is as in **A**. **(D)** Plots of raw and BBUM-FDR-adjusted *p* values. Colors are as in **A**. The *p*_*BBUM*_ significance cutoff at 0.05 is shown as orange dashed lines. Five data points for extremely small *p* and *p*_*BBUM*_ values that fell out of the range to the right are not shown in the plot for contact-inhibited MEF cells: miR-7a-5p, miR-335-3p, miR-376b-3p, miR-154-3p, and miR-672-5p.

Three other edge cases proposed to be ZSWIM8 substrates by the previous method were not identified at *p*_*BBUM*_ < 0.05 when using the BBUM model (Fig. 3A, Table S1). One of these was miR-7-5p in clonal ZSWIM8 knockout cells. This known TDMD substrate [15] was not sufficiently upregulated in knockout K562 cells to reach significance over the relatively high background variability of this dataset (Fig. 3A). Nonetheless, statistical significance was readily achieved for miR-7-5p in K562 cells when using datasets in which ZSWIM8 was knocked down using CRISPRi, which led to much lower background variability than observed with clonal knockout cells (Fig. 3A). Likewise, in a dataset analyzing a different ZSWIM8-knockout line (Han et al., 2020), miR-7-5p upregulation met the significance threshold using the BBUM approach (Fig. 3C).

Thus, on the whole, using the BBUM model, candidate primary substrates of ZSWIM8 were identified while implementing a consistent and predetermined FDR confidence value across all cellular contexts examined, without noticeably sacrificing the apparent sensitivity or specificity of the previous approach. We attribute this success to the ability of the BBUM model to adjust the varying spread of background *p* values to a consistent range (Fig. 3D).

### BBUM correction applies a consistent statistical stringency

To benchmark the performance of our approach, we randomly generated simulated datasets of *p* values containing varying levels of primary and background secondary signal under the BBUM distribution. We compared the empirical FDR of our BBUM strategy, using *p*_*BBUM*_ < 0.05 as the significance cutoff, against that of the method used previously by Shi et al. [9]. The BBUM approach had a mean FDR of 0.050 ± 0.002 (95% confidence interval (CI)) (Fig. 4A). The Shi et al. method produced a comparable mean FDR of 0.036 ± 0.002 but was less constant, as measured by the coefficient of variation (CV) of the FDR, which was 2.47 ± 0.11 for the previous method, compared to 1.55 ± 0.08 for the BBUM method. Thus, BBUM correction produced an accurate and consistent FDR when evaluated using simulated datasets.

**Figure 4.**
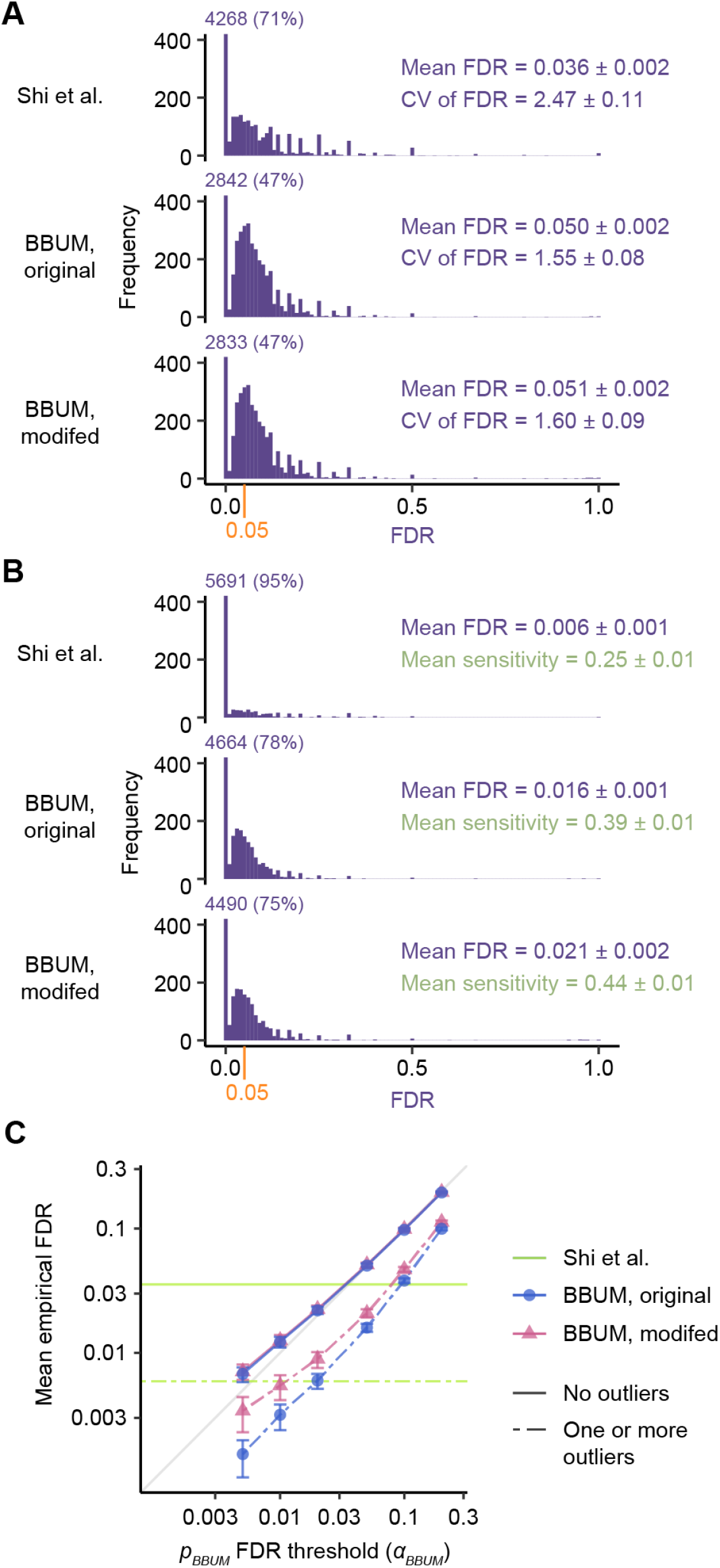
Benchmarking of correction procedures and significance-testing methods. **(A)** Histograms of empirical FDR for 6000 simulated *p* value datasets with no outliers. Results are plotted for the extreme-value method used by Shi et al., the original BBUM procedure without outlier trimming at *p*_*BBUM*_ < 0.05, and the modified BBUM procedure with automatic outlier trimming at *p*_*BBUM*_ < 0.05. The frequency of the density near 0, which exceeded the *y*-axis scale, is indicated alongside its fraction among all trials for each histogram. The FDR cutoff of the BBUM methods at 0.05 is indicated on the x-axis in orange. The mean FDR of all simulation trials is indicated on the right with corresponding 95% CIs by bootstrapping. The CV of FDR is similarly indicated. **(B)** As in **A**, but for 6000 simulations with at least one programmed outlier. The mean FDR and mean sensitivities are indicated with corresponding 95% CIs by bootstrapping. **(C)** The mean empirical FDR of significance-testing methods at different *p*_*BBUM*_ significance thresholds. Results are plotted for all six threshold values tested. If the empirical FDR matched the predetermined FDR cutoff, the point would lie on the gray diagonal line. Error bars indicate 95% CIs by bootstrapping. Because the Shi et al. method does not allow quantitative adjustments of significance thresholds, results are shown as green horizontal lines with the mean empirical FDR values as the *y*-intercepts.

BUM models are reported to be sensitive to outliers with extremely small *p* values due to the asymptotic behavior at zero of the likelihood function of the type of beta distribution used [13]. Indeed, we found that adding artificial extreme outliers to downregulated miRNAs in either empirical or simulated datasets could cause the BBUM procedure to overcorrect for secondary effects. The previous approach by Shi et al. [9] was even more prone to overcorrection, with a single extreme outlier able to cause all of the upregulated miRNAs to be designated as background. The influence of outliers was also illustrated in the Han et al. [10] dataset for K562 cells. Two extreme outliers within this dataset prevented any miRNAs to be designated as primary ZSWIM8 substrates when using the approach of Shi et al. [9] and severely weakened the significance of miR-7-5p when using the BBUM model (Fig. 3C).

To mitigate the effects of outliers, we developed a conservative outlier detection method that used the fitted *r* parameter of the BBUM model to identify and trim probable outliers from downregulated data points. The performance of the modified BBUM procedure was similarly benchmarked using simulated *p-*value datasets, with and without added outliers. In datasets without added outliers, the modified BBUM procedure did not have significantly increased mean FDR (0.051 ± 0.002) or CV of FDR (1.60 ± 0.09) when compared to the original BBUM procedure (Fig. 4B). In datasets with added outliers, the modified BBUM procedure was somewhat improved over the unmodified procedure, with mean FDR increasing from 0.016 ± 0.001 for the original BBUM procedure to 0.021 ± 0.002 for the modified procedure and mean sensitivity increasing from 0.39 ± 0.01 to 0.44 ± 0.01 (Fig. 4B). Importantly, both the original and the modified BBUM procedures were less sensitive to outliers than the previous method of Shi et al. [9], which had a mean FDR of 0.006 ± 0.001 and mean sensitivity of 0.25 ± 0.01 in the presence of one or more outliers (Fig. 4B). Moreover, the modified BBUM procedure successfully identified and trimmed the two outliers in the K562 dataset (Fig. 3C), as well as any artificial outliers we added to other empirical datasets.

The ad hoc method by Shi et al. [9] provides a fixed stringency for each dataset. In contrast, the BBUM method allows the stringency to be quantitatively tuned by choosing the significance cutoff for *p*_*BBUM*_ to suit the needs of the experiment or hypothesis at hand. Therefore, we extended our benchmarking analysis to a range of possible significance cutoffs for *p*_*BBUM*_, and assessed the accuracy of the BBUM method at each cutoff *α*_*BBUM*_ in our simulations. When no outliers were present, both the original and the modified BBUM methods faithfully achieved results at the predetermined FDR at all *α*_*BBUM*_ values tested (Fig. 4C). When one or more outliers were present, the modified BBUM method partially mitigated the overcorrection of secondary effects by the original BBUM method at all *α*_*BBUM*_ values tested, especially when *α*_*BBUM*_ was small, which was the condition in which overcorrection was most severe in the original BBUM method (Fig. 4C). In fact, the overcorrection of the modified BBUM procedure at all *α*_*BBUM*_ values tested was no worse than that seen at *α*_*BBUM*_ = 0.05, where the mean empirical FDR was 0.021 ± 0.002 (Fig. 4B, C). Hence, the BBUM procedure performed robustly in both ideal and non-ideal scenarios and performed significance testing with improved consistency and flexibility, as well as lower sensitivity to outliers.

## DISCUSSION

Our BBUM method will help to more rigorously identify the miRNAs subject to TDMD. We suspect that this approach will also be useful in other analyses in which the primary effect of a regulatory phenomenon causes changes in one direction, whereas secondary effects and background variability cause changes in either direction. For example, the BBUM approach might be useful for analyses of miRNA regulation, wherein primary targets are expected to only increase upon deletion or knockdown of the miRNA. Other potential uses include mRNA analyses identifying the targets of transcriptional inhibitors or proteomic analyses identifying the targets of ubiquitin ligases or other degradation phenomena.

## METHODS

### Analysis of small-RNA sequencing results

Raw read counts of miRNAs from small-RNA sequencing data were obtained from previously published datasets, and filtered and normalized as described [9]. Briefly, sequencing reads were matched to the first 19 nt of mature miRNA sequences downloaded from TargetScan7 [16] and miRNAs were filtered to have at least five matched reads in each sequencing library. Mean fold changes and *p* values were then calculated using DESeq2 (v1.32.0) [1], using the default Wald test method, without the lfcShrink() function, and with independentFiltering = FALSE in results().

### Significance testing by the Shi et al. method

To call changes as significant by the method of Shi et al., default FDR-adjusted *p* values calculated using the Benjamini-Hochberg procedure from DESeq2 were used. The *p*_*adj*_ threshold for significance was chosen as the most permissive value out of a defined sequence of canonical critical values (0.05, 0.01, 0.001, 0.0001, …) that excluded all downregulated miRNAs in the dataset. For example, in iMNs, the strongest downregulated miRNA had a *p*_*adj*_ value of 0.000262, and thus the threshold was adjusted to 0.0001.

### The BBUM statistical model of *p* values

The *p* values from the DE experiments can be reasonably modeled as a random variable *X* following a BBUM distribution:

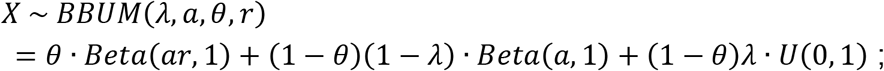

the probability density function (PDF) of the model is defined as:

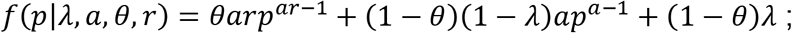

likewise, the cumulative distribution function (CDF) is defined as:

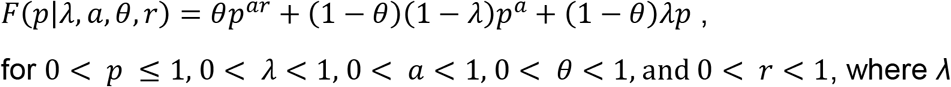

for 0 < *p* ≤ 1, 0 < *λ* < 1, 0 < *a* < 1, 0 < *θ* < 1, and 0 < *r* < 1, where *λ* represents the fraction of null distribution density over all density except primary signal, *θ* represents the fraction of primary signal distribution density over all density, *a* describes the shape of the peak of secondary signal, and *r* describes the ratio between the shape parameters of the primary and secondary signal components. The PDF asymptotes to infinity as *p* approaches 0, and monotonically decreases as *p* goes from 0 to 1.

### BBUM model fitting and parameter estimation

Given a set of *p* values ***p***, the BBUM distribution model was fit to ***p*** using a modified maximum likelihood estimation (MLE) method. While varying a shared set of parameters, *p* values associated with upregulated miRNAs, ***p***+, were fit to the full BBUM function, whereas *p* values associated with downregulated miRNAs, ***p***−, were fit to the same BBUM function with *θ* fixed at 0 and *r* fixed at 1, which corresponded to a zero component for primary effects (Fig. 1D). The total log-likelihood *ℓ* was used as the maximization target for MLE fitting and was defined as the sum of log-likelihood values of the two halves:

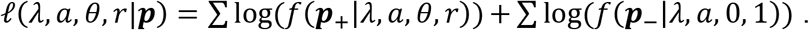

Fitting was performed using the optim() function in R using the default Broyden-Fletcher-Goldfarb-Shanno (BFGS) algorithm for a maximum of 200 iterations. Unless otherwise stated, parameters *λ*, *a*, and *r* were bounded to (0, 1). *θ* was bounded to (0, 1 – 2*α*_*BBUM*_), where *α*_*BBUM*_ was the critical threshold of BBUM-adjusted *p* values used for significance testing, to prevent the local minimum near *θ* = 1 where all upregulated points would be regarded as primary signal when there is very low or no primary signal.

The four parameters were bounded by transforming the values using the logit (log-odds) function. Given a parameter *x* with lower bound *x*_*lb*_ and upper bound *x*_*ub*_, the value was transformed as:

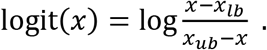

For each dataset, fitting was initiated at each of six sets of fixed initial parameter values (Table S2). The initial values for *θ* were linearly rescaled to its bounds. Out of the six attempts, the successfully converged solution with the highest total log-likelihood was chosen as the final solution.

### Outlier detection and trimming

Due to the asymptotic behavior of the likelihood function at zero, BBUM model fitting can be prone to overcorrection for secondary effects when outliers with extremely low *p* values among downregulated miRNAs are present. To mitigate this, we developed and implemented a conservative method for outlier detection and trimming. The data were first preliminarily fit to the model with a wider bound for *r* at (0, 10). When very strong outliers were present among downregulated miRNAs, the algorithm would converge to a value of *r* > 1, which implied a stronger secondary effect than primary effect, violating the assumption of our model and suggesting the existence of unexpectedly strong signal in the background. An increasing number of downregulated miRNAs with the strongest *p* values was then iteratively trimmed as outliers until the algorithm converged to a value of *r* < 1, unless the condition was not met after trimming 5% or 10 downregulated miRNAs, whichever was lower.

### Significance testing

While only considering upregulated data points, the expected FDR level of falsely calling either null points or secondary signal as primary signal can be calculated at any given cutoff *α* for raw *p* values. We employed a strategy for adjusting *p* values that resembled the one that DESeq2 adopts for the Benjamini-Hochberg procedure [1]. For every raw *p* value of an upregulated miRNA, we calculated the expected FDR value at that value using the BBUM model and denoted it as the BBUM-FDR-adjusted *p* value (*p*_*BBUM*_) for significance testing. Thus, to control for FDR at a pre-determined cutoff, such as *α*_*BBUM*_ = 0.05, all points with *p*_*BBUM*_ < 0.05 would be called as significant. The expected FDR was calculated as

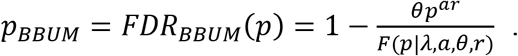

### Simulation of *p* values and benchmarking

To benchmark strategies for significance calling and correction, DE *p* values with primary and secondary effects were simulated using the BBUM model. For each simulation, the total number of *p* values, *n*, was generated from a uniform distribution, and the number of upregulated points was determined through a binomial distribution:

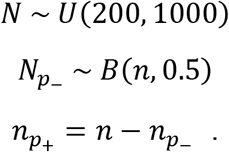

The values of *p* were then modeled as a random variable *X*, which followed a compound distribution of the upregulated and downregulated halves, *X*_*p+*_ and *X*_*p−*_, respectively:

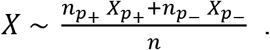

For each half of the dataset, *p* values were simulated under respective BBUM models. If programmed outliers were simulated among downregulated points, the outliers were simulated as a separate beta component similarly as the primary signal, where

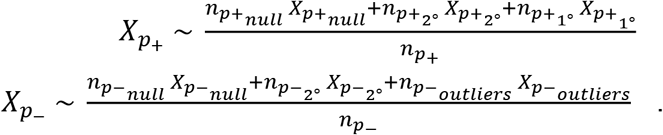

The proportions of the mixture components were drawn from binomial distributions using the values of *λ*, *θ*, and *θ’*, where *θ’* represented the *θ* parameter for outliers, where:

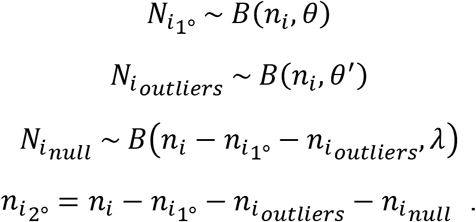

Each component of the BBUM models was modeled as previously described, where

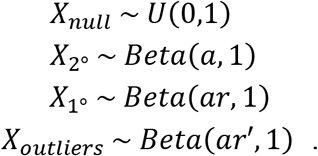

For each simulation. the BBUM parameters were randomly generated from uniform or exponential distributions over reasonable expected ranges of values:

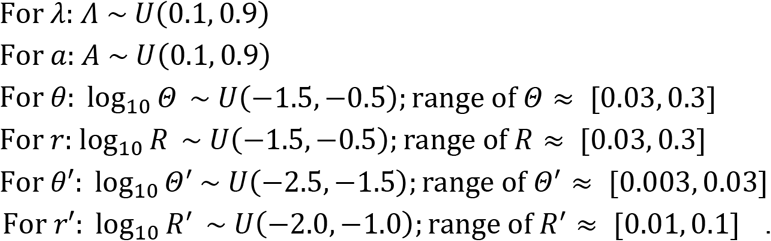

Only simulations with at least three data points under primary effects were accepted, to allow sufficient true hit data points for benchmarking. For simulations with outliers, only simulations with at least one outlier were accepted.

Benchmarking statistics, such as FDR and sensitivity, were calculated by comparing the points called as significant with the BBUM distribution components they were drawn from, using the following equations (Fig. 1C):

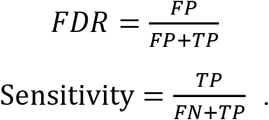

For each simulation, 6000 simulated datasets were generated. Confidence intervals for mean and CV statistics were generated by ordinary bootstrapping using the boot package in R. Datasets were resampled 3000 times, and nonparametric 95% confidence intervals were defined by the empirical bootstrap confidence intervals, using the “basic” method of the boot.ci function. Based on the central limit theorem, confidence intervals were presented as the mean deviation of the lower and upper intervals from the sample mean.

### Artificial outliers for datasets

To assess the potential impact of outliers on different adjustment methods using empirical datasets from Shi et al. [9], we added to each dataset one to five extreme downregulated outliers with raw *p* values at 10^−300^.

## ABBREVIATIONS

TDMD: Target-mediated microRNA degradation
BBUM: Bi-beta uniform mixture
FDR: False discovery rate
DE: Differential expression
CI: Confidence interval

## DECLARATIONS

### Ethics approval and consent to participate

Not applicable.

### Consent for publication

Not applicable.

### Availability of data and materials

All data analyzed during this study are included in these published articles [9, 10] and their supplementary information files.

The BBUM fitting and significance-calling algorithm is available as an R package at https://github.com/wyppeter/bbum. Other code used for this work, including data analyses, simulations, and data visualization, is available at https://github.com/wyppeter/BBUM-TDMD_2022.

### Authors’ contributions

P.Y.W. conceived the approach, designed and implemented the model, and analyzed the data, with input from D.P.B. Both authors wrote the manuscript.

## Acknowledgements

We thank E. R. Kingston, C. Y. Shi, and others in the Bartel laboratory for helpful discussions and feedback.

## ADDITIONAL FILES

**Additional file 1**.

Table S1: miRNAs in the Shi et al. datasets identified as ZSWIM8-sensitive.

Table S2: Raw initial parameter values for BBUM model fitting.

