## Additional File 1 for "A statistical approach for identifying primary substrates of ZSWIM8-mediated microRNA degradation in small-RNA sequencing data"

**Table S1.** miRNAs in the Shi et al. datasets identified as ZSWIM8-sensitive.

| Cell line | miRNA | Notes |
|---|---|---|
| ***Not changed*** | | |
| S2 | dme-miR-7-5p | |
| A549 | hsa-miR-7-5p | |
| HeLa | hsa-miR-7-5p | |
| K562 (KD, A) | hsa-miR-7-5p | |
| K562 (KD, B) | hsa-miR-7-5p | |
| MCF-7 | hsa-miR-7-5p | |
| iMN | mmu-miR-7a-5p | |
| MEF | mmu-miR-7a-5p | |
| iMN | mmu-miR-7b-5p | |
| S2 | dme-miR-9b-5p | |
| S2 | dme-miR-9c-5p | |
| S2 | dme-miR-12-5p | |
| MEF | mmu-miR-17-5p | |
| MEF | mmu-miR-20a-5p | |
| HeLa | hsa-miR-29b-3p | |
| iMN | mmu-miR-29b-3p | |
| MEF | mmu-miR-29b-3p | |
| iMN | mmu-miR-33-5p | |
| MEF | mmu-miR-33-5p | |
| MEF | mmu-miR-92a-3p | |
| MEF | mmu-miR-93-5p | |
| MCF-7 | hsa-miR-145-5p | |
| A549 | hsa-miR-154-3p | |
| HeLa | hsa-miR-154-3p | |
| iMN | mmu-miR-154-3p | |
| MEF | mmu-miR-154-3p | |
| MEF | mmu-miR-181b-5p | |
| S2 | dme-miR-190-5p | |
| MEF | mmu-miR-193a-3p | |
| MEF | mmu-miR-195a-5p | |
| S2 | dme-miR-277-3p | |
| S2 | dme-miR-279-3p | |
| MEF | mmu-miR-322-5p | |
| HeLa | hsa-miR-335-3p | |
| MCF-7 | hsa-miR-335-3p | |
| iMN | mmu-miR-335-3p | |
| MEF | mmu-miR-335-3p | |
| iMN | mmu-miR-341-3p | |
| A549 | hsa-miR-376a-3p | |

| | | |
|---|---|---|
| HeLa | hsa-miR-376a-3p | |
| MCF-7 | hsa-miR-376b-3p | |
| iMN | mmu-miR-376b-3p | |
| MEF | mmu-miR-376b-3p | |
| iMN | mmu-miR-409-3p | |
| MEF | mmu-miR-425-5p | |
| iMN | mmu-miR-431-5p | |
| MEF | mmu-miR-450b-5p | |
| MEF | mmu-miR-485-3p | |
| iMN | mmu-miR-495-3p | |
| MEF | mmu-miR-495-3p | |
| MEF | mmu-miR-503-5p | |
| MEF | mmu-miR-532-5p | |
| A549 | hsa-miR-543-3p | |
| HeLa | hsa-miR-543-3p | |
| iMN | mmu-miR-543-3p | |
| MEF | mmu-miR-543-3p | |
| MCF-7 | hsa-miR-652-5p | |
| iMN | mmu-miR-665-3p | |
| MEF | mmu-miR-665-3p | |
| iMN | mmu-miR-672-5p | |
| MEF | mmu-miR-672-5p | |
| iMN | mmu-miR-744-5p | |
| S2 | dme-miR-996-3p | |
| S2 | dme-miR-999-3p | |
| S2 | dme-miR-1002-5p | |
| S2 | dme-miR-1012-3p | |
| MCF-7 | hsa-miR-1247-5p | |
| MEF | mmu-miR-1247-5p | |
| ***Added*** | | |
| MCF-7 | hsa-miR-20a-5p | Also found in MEF. |
| MEF | mmu-miR-193b-3p | Family member also found in MEF. |
| MEF | mmu-miR-369-3p | |
| MEF | mmu-miR-409-3p | Also found in iMN. |
| ***Removed*** | | |
| K562 (KO) | hsa-miR-7-5p | Also found in multiple other systems. |
| iMN | mmu-miR-92a-3p | Also found in MEF. |
| iMN | mmu-miR-297c-5p | |

**Table S2.** Raw initial parameter values for BBUM model fitting.

| Initial value set | $\lambda$ | $a$ | $\theta$ | $r$ |
|---|---|---|---|---|
| 1 | 0.9 | 0.9 | 0.1 | 0.1 |
| 2 | 0.9 | 0.1 | 0.9 | 0.1 |
| 3 | 0.9 | 0.1 | 0.1 | 0.9 |
| 4 | 0.1 | 0.1 | 0.9 | 0.9 |
| 5 | 0.1 | 0.9 | 0.1 | 0.9 |
| 6 | 0.1 | 0.9 | 0.9 | 0.1 |
